# Comparison of Cepheid Xpert Xpress and Abbott ID Now to Roche cobas for the Rapid Detection of SARS-CoV-2

**DOI:** 10.1101/2020.04.22.055327

**Authors:** Marie C. Smithgall, Ioana Scherberkova, Susan Whittier, Daniel A. Green

## Abstract

The SARS-CoV-2 pandemic has created an urgent and unprecedented need for rapid large-scale diagnostic testing to inform timely patient management. This study compared two recently-authorized rapid tests, Cepheid Xpert Xpress SARS-CoV-2 and Abbott ID Now SARS-CoV-2 to the Roche cobas SARS-CoV-2 assay. A total of 113 nasopharyngeal swabs were tested, including 88 positives spanning the full range of observed C_t_ values on the cobas assay. Compared to cobas, the overall positive agreement was 73.9% with ID Now and 98.9% with Xpert. Negative agreement was 100% and 92.0% for ID Now and Xpert, respectively. Both ID Now and Xpert showed 100% positive agreement for medium and high viral concentrations (C_t_ value <30). However, for C_t_ values >30, positive agreement was 34.3% for ID Now and 97.1% for Xpert. These findings highlight an important limitation of ID Now for specimens collected in viral or universal transport media with low viral concentrations. Further studies are needed to evaluate the performance of ID Now for direct swabs.

## Introduction

Severe acute respiratory virus coronavirus 2 (SARS-CoV-2) emerged in Wuhan, China in December 2019 and has since rapidly spread across the world, causing a global pandemic of coronavirus disease (COVID-19). The majority of cases are mild to moderate, but severe infections have overwhelmed healthcare systems in the United States, particularly in New York City. Real-time polymerase chain reaction (RT-PCR) of viral RNA from nasal or nasopharyngeal swabs has become the standard method used to confirm diagnosis. The first quantitative RT-PCR test for detecting SARS-CoV2 was designed and distributed in January 2020 by the World Health Organization (WHO) (1). In the United States and many other countries, however, the slow rollout of large-scale diagnostic testing and the long turnaround times associated with laboratory tests, particularly those sent to reference laboratories, have significantly hampered public health efforts to contain the outbreak.

In contrast, commercially-available rapid diagnostic assays can better inform timely patient management decisions to guide the need for quarantine, isolation, contact tracing, and therapeutic management. Beginning in March 2020, multiple SARS-CoV-2 rapid tests received Emergency Use Authorization (EUA) from the U.S. Food and Drug Administration (FDA). However, manufacturer submissions for EUA only require evaluation of the limit of detection and cross-reactivity of their assays, and do not address other important performance characteristics such as accuracy, precision, and reproducibility. In addition, most manufacturer submissions include studies of contrived positive samples with spiked viral RNA, and do not assess performance on clinical patient specimens. Two recently-authorized rapid tests, Xpert Xpress SARS-CoV-2 (Cepheid, Sunnyvale, CA) and ID Now SARS-CoV-2 (Abbott, Chicago, IL) have been manufactured at wide-scale and distributed to numerous medical centers around the country. While limited studies on these two assays have been recently published, the number of patient samples evaluated to date has been relatively small, and significant questions remain about the accuracy of these tests across the full spectrum of viral loads (2–4). Utilizing the high volume of patient testing performed at our medical center in New York City, we sought to evaluate and compare the performance of these two rapid assays across a range of clinical samples.

## Materials and Methods

### Study Design and Data Analysis

Deidentified remnant patient samples that underwent routine clinical testing with the cobas SARS-CoV-2 assay on the 6800 platform (Roche Diagnostics, Indianapolis, IN) were used to evaluate the Xpert and ID Now assays. Residual nasopharyngeal (NP) swabs in transport media were held at 4° C prior to testing on the Xpert and ID NOW platforms, with all testing completed within 48 hours of sample collection. Testing was performed according to the manufacturers’ instructions on two separate ID Now instruments and a single GeneXpert Infinity instrument.

A total of 113 NP swabs collected in 3 mL of viral transport media (M4RT VTM; ThermoFisher Scientific, Waltham, MA) or universal transport media (UTM; Becton Dickinson and Co., Franklin Lakes, NJ) were included. The specimens were collected from April 8 to April 13, 2020 and included 111 adult and 2 pediatric patients who were all seen in inpatient or emergency room locations.

To evaluate assay performance at varying viral concentrations, 88 positive specimens were selected to represent the full range of observed C_t_ values on the cobas assay, ranging from 14 – 38 cycles. Positive agreement and 95% confidence intervals for the Xpert and ID Now assays were calculated using cobas as the reference test. An additional 25 negative specimens were selected to evaluate negative agreement.

### Assay Descriptions

The cobas assay utilizes RT-PCR to amplify and detect two viral targets: ORF1 a/b, a non-structural region that is unique to SARS-CoV-2 and a conserved region in the E-gene, which is a structural protein envelope for pan-Sarbecovirus detection. The Xpert assay is an automated RT-PCR that amplifies and detects two viral targets: N2, a nucleocapsid recombinant protein unique to SARS-CoV-2 and a region in the structural envelope E-gene. The ID Now assay uses proprietary isothermal nucleic acid amplification technology for qualitative detection of SARS-CoV-2 RdRp gene using fluorescent reporter probes.

This study was approved by the Columbia University Irving Medical Center Institutional Review Board.

## Results

Of the 113 patient specimens, 111 were from adults ranging in age from 23 to 101 years and two were from pediatric patients, aged 1 day and 5 days old. The average age was 65 years for positive samples and 43 years for negative samples. Overall, the majority of positive samples were from males (60.2%) and negative samples from females (68.0%) (Table 1).

**Table 1:**
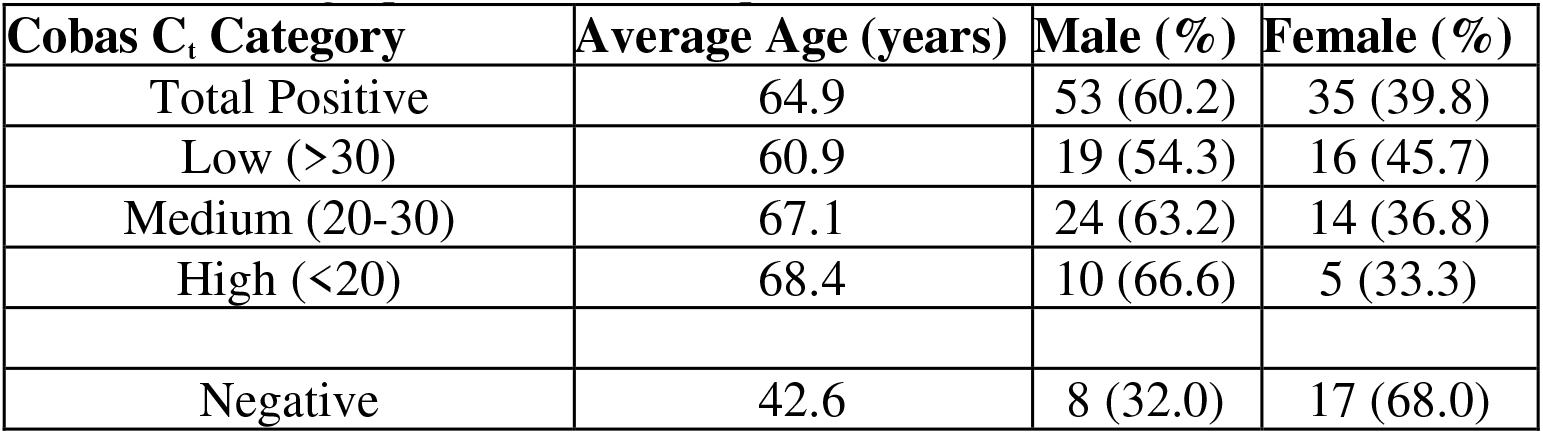
Demographics of included patients

Testing results by Abbott ID Now and Cepheid Xpert are shown in Table 2 and Figure 1. Compared to cobas, the overall positive agreement with ID Now was 73.9% (95% Confidence Interval (CI): 63.2 – 82.3%) and with Xpert was 98.9% (95% CI 92.9 – 100%). Negative agreement was 100% (95% CI 83.4 – 100%) and 92.0% (95% CI 72.4 – 98.6%) for ID Now and Xpert, respectively. Both ID Now and Xpert showed 100% positive agreement for medium and high viral concentrations, defined as C_t_ value <30. However, for C_t_ values >30, positive agreement for ID Now was 34.3% (95% CI 19.7 – 52.2%), whereas it was 97.1% (95% CI 83.4 – 99.8%) for Xpert. Notably, one sample detected by Xpert was a presumptive positive based on detection of the E-gene target but not the N2 target. There were also two samples that tested negative by cobas but positive by the Xpert. These samples had C_t_ values >40 for the N2 target only without detection of the E-gene target. For the E-gene target, C_t_ values were generally 1 cycle lower for Xpert than cobas (Supplemental Materials, Table S1 and Figure S1).

**Figure 1A:**
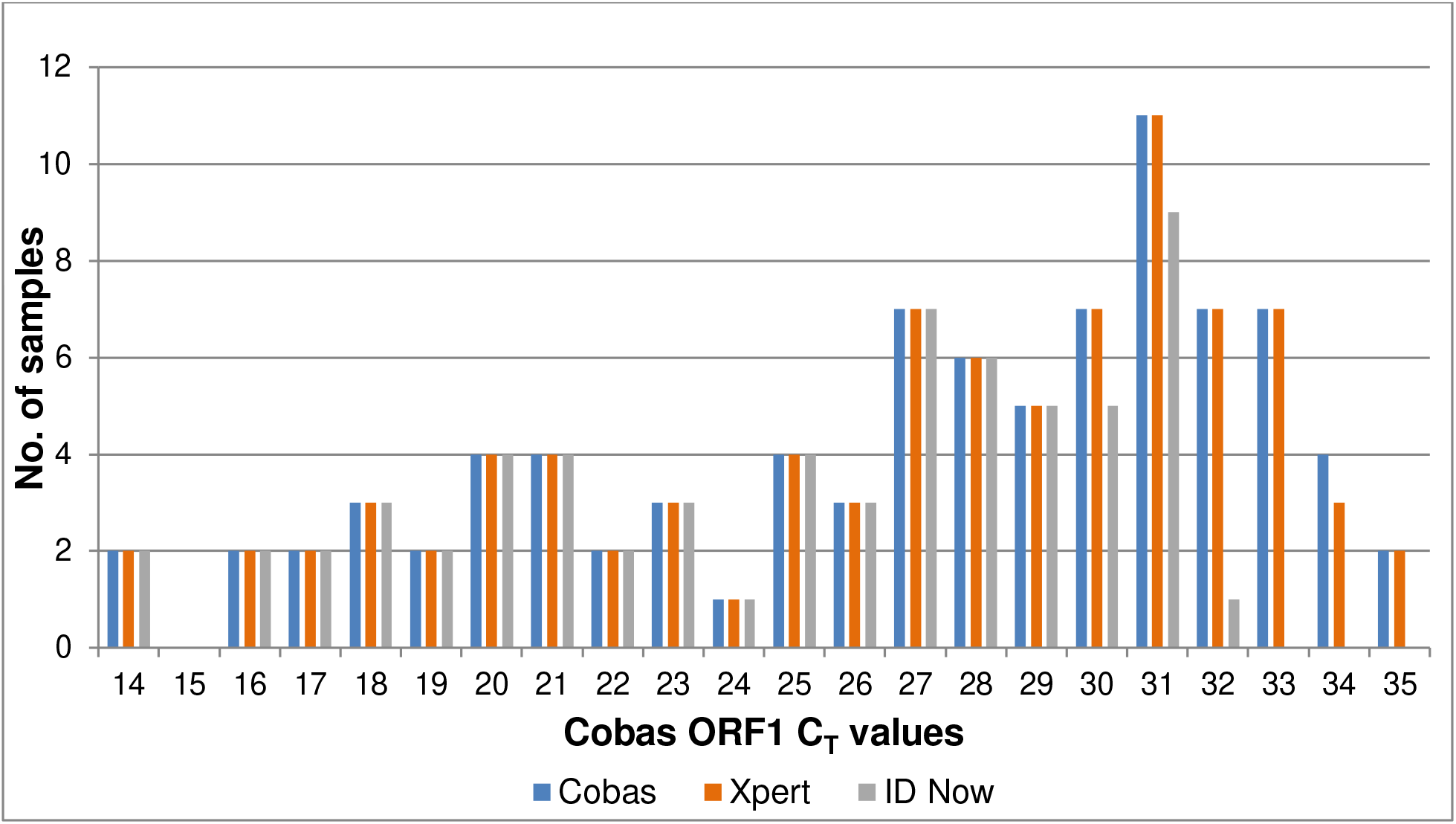
Frequency distribution of cycle threshold (C_t_) values for positive patient samples by Roche cobas SARS-CoV-2 (blue), Cepheid Xpert Xpress SARS-CoV-2 (orange) and Abbott ID Now SARS-CoV-2(gray) assays. Roche cobas Target 1 (ORF1) C_t_ values were rounded to the nearest whole number.

**Figure 1B:**
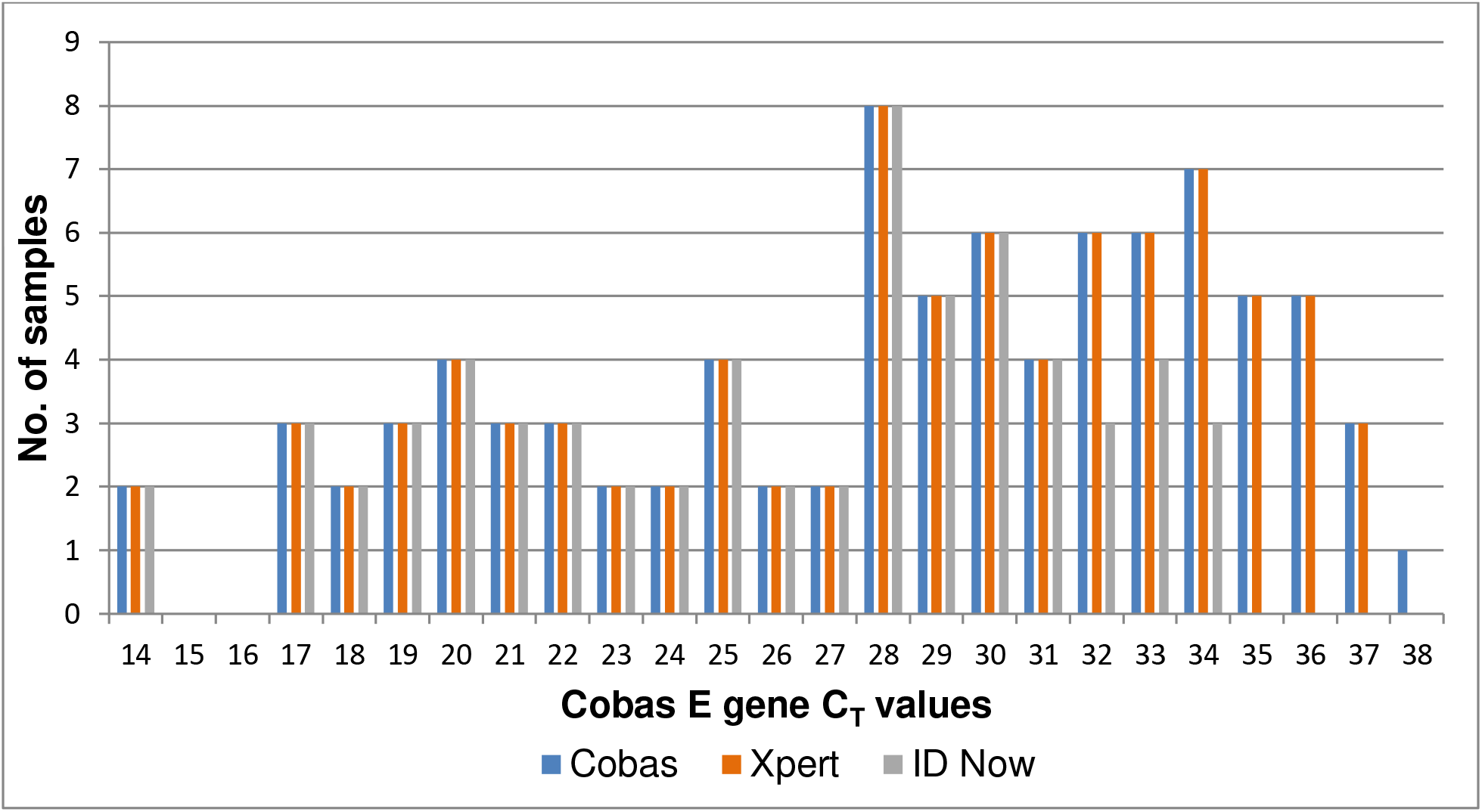
Frequency distribution of cycle threshold (C_t_) values for positive patient samples by Roche cobas SARS-CoV-2 (blue), Cepheid Xpert Xpress SARS-CoV-2 (orange) and Abbott ID Now SARS-CoV-2(gray) assays. Roche cobas Target 1 (ORF1) C_t_ values were rounded to the nearest whole number.

**Table 2:**
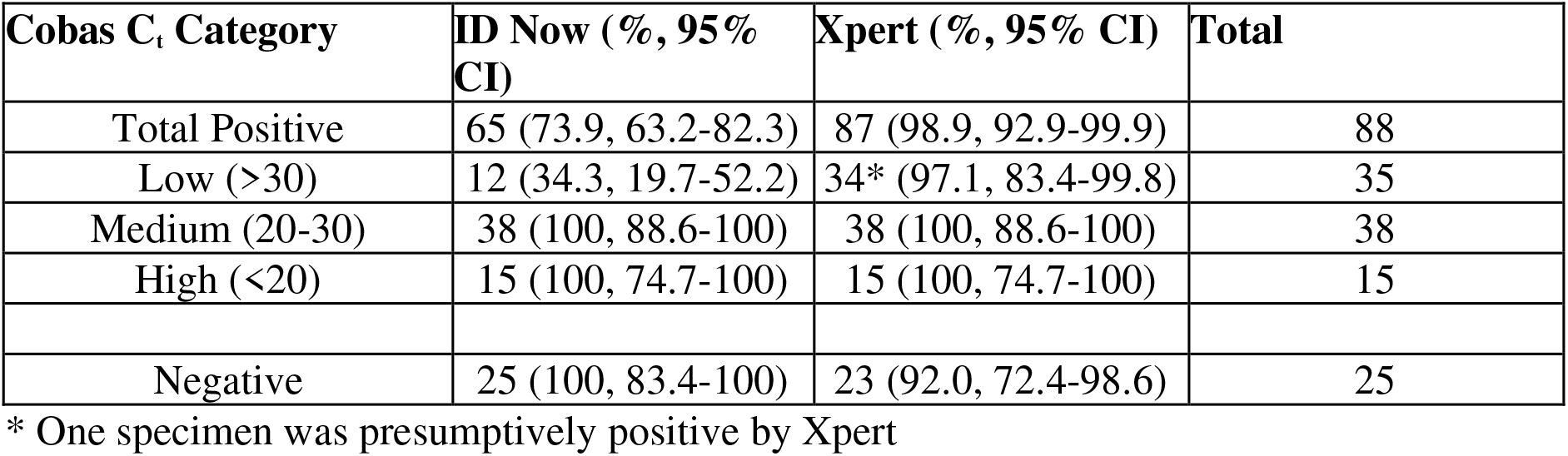
Positive and negative agreement of Abbott ID Now SARS-CoV-2 and Cepheid Xpert Xpress SARS-CoV-2 with Roche cobas SARS-CoV-2

| **Cobas C<sub>t</sub> Category** | **ID Now (% , 95% CI)** | **Xpert (% , 95% CI)** | **Total** |
| --- | --- | --- | --- |
| Total Positive | 65 (73.9, 63.2-82.3) | 87 (98.9, 92.9-99.9) | 88 |
| Low (>30) | 12 (34.3, 19.7-52.2) | 34* (97.1, 83.4-99.8) | 35 |
| Medium (20-30) | 38 (100, 88.6-100) | 38 (100, 88.6-100) | 38 |
| High (<20) | 15 (100, 74.7-100) | 15 (100, 74.7-100) | 15 |
| Negative | 25 (100, 83.4-100) | 23 (92.0, 72.4-98.6) | 25 |

\* One specimen was presumptively positive by Xpert

## Discussion

To meet the urgent need for wide-scale diagnostic testing during the COVID-19 pandemic, multiple rapid molecular tests have recently been authorized by the US FDA, some of which are available in point-of-care (POC) or near patient settings. However, very few studies have been published to date on the relative performance characteristics of these assays, especially for patient specimens representing a wide range of viral concentrations (2, 3).

In this comparative analysis, the Xpert assay showed a very high level of agreement with the cobas assay across the entire range of tested C_t_ values, including low-level positives. These findings confirm those published by Moran *et al.* (2) and show a high level of agreement between these two assays using an expanded number of positive clinical samples. In contrast, the ID Now assay reliably detected specimens with C_t_ values ≤30, but did not detect the majority of specimens with C_t_ values ≥ 30. Whereas Rhoades *et al.* (3) found an overall high level of agreement between ID Now and the modified CDC assay, our findings highlight an important limitation of the ID Now for low-level positives. While both studies evaluated nasopharyngeal swabs eluted in transport media, it is important to note that the EUA for ID NOW was recently updated to remove the indication for swabs in transport media (5). Our data support that the EUA was appropriately modified, as samples may become too dilute in VTM and low-level positives may falsely test negative.

In contrast to batch testing and the higher complexity required for the cobas assay, both Xpert and ID Now offer shorter turnaround times and availability in near-patient settings. However, the two assays differ in throughput capacity and run time. Each ID Now platform can run only a single specimen at a time, with results available in 13 minutes or less. Xpert can be run on larger, random-access platforms that allow for significantly higher throughput, with results available in 45 minutes. Both assays are available for use in POC settings, which introduces both benefits and drawbacks. On the one hand, POC molecular testing delivers the shortest possible interval between sample collection and result, which can facilitate faster clinical decision-making. However, concerns related to assay performance, quality management, and safety in the POC setting still remain. Studies of POC molecular testing for influenza and respiratory syncytial virus have shown promising results, but also highlight some of these concerns (6–9). The risk of contamination and false positives is also much higher when testing is performed outside of a controlled environment and by non-laboratory trained personnel. Nevertheless, the unique circumstances of a rapidly-spreading pandemic may ultimately necessitate the widescale adoption of POC assays in completely new patient settings, such as walk up or drive-thru testing centers.

Limitations of this study include the relatively few number of samples from pediatric patients, as only two samples from patients aged 1 day and 5 days were included, and both of these were negative on all three testing methods. We were also only able to evaluate ID Now for specimens in transport media. The performance of this assay with direct nasal swabs requires further evaluation in subsequent studies. Another limitation is the use of the cobas assay as the comparator assay. Two samples that were identified as positive only by Xpert on the basis of N2 nucleocapsid gene detection were negative for both targets on cobas. Whether these samples were truly positive or truly negative could not be determined.

Fast, readily available, and reliable test results are critically important during this pandemic, and each of the three assays evaluated in this study holds promise to deliver valuable clinical information. Further head-to-head comparisons of molecular tests will be important in order to establish the usefulness of each method and to help medical providers determine the most appropriate diagnostic tests to best serve their communities.

## References

1. Sheridan C. Coronavirus and the race to distribute reliable diagnostics. Nature Biotechnology 38, 382–384 (2020) doi: 10.1038/d41587-020-00002-2.

2. Moran A, Beavis KG, Matushek SM, et al. The Detection of SARS-CoV-2 using the Cepheid Xpert Xpress SARS-CoV-2 and Roche cobas SARS-CoV-2 Assays. J Clin Microbiol. 2020 Apr 17. pii: JCM.00772-20. doi: 10.1128/JCM.00772-20

3. Rhoads D, Cherian S, Roman K et al. Comparison of Abbott ID Now, Diasorin Simplexa, and CDC FDA EUA methods for the detection of SARS-CoV-2 from nasopharyngeal and nasal swabs from individuals diagnosed with COVID-19. J Clin Microbiol. 2020 Apr 17. pii: JCM.00760-20. doi: 10.1128/JCM.00760-20

4. Poljak M, Korva M, Gašper NK, et al. Clinical evaluation of the cobas SARS-CoV-2 test and a diagnostic platform switch during 48 hours in the midst of the COVID-19 pandemic. Journal of Clinical Microbiology Apr 2020, JCM.00599-20; DOI: 10.1128/JCM.00599-20.

5. ID NOW COVID-19 Technical Brief and Sample Type Labeling Update. April 2020.

6. Hazelton B, Gray T, Ho J, et al. Detection of influenza A and B with the Alere ™ i Influenza A & B: a novel isothermal nucleic acid amplification assay. Influenza Other Respir Viruses. 2015;9(3):151–154. doi:10.1111/irv.12303.

7. Chen JH, Lam HY, Yip CC et al. Evaluation of the molecular Xpert Xpress Flu/RSV assay vs. Alere i Influenza A & B assay for rapid detection of influenza viruses. Diagn Microbiol Infect Dis. 2018 Mar;90(3):177–180. doi: 10.1016/j.diagmicrobio.2017.11.010.

8. Hassan F, Hays LM, Bonner A, et al. Multicenter Clinical Evaluation of the Alere i Respiratory Syncytial Virus Isothermal Nucleic Acid Amplification Assay. J Clin Microbiol. 2018;56(3):e01777–17. Published 2018 Feb 22. doi:10.1128/JCM.01777-17.

9. Nolte FS, Gauld L, Barrett SB. Direct Comparison of Alere i and cobas Liat Influenza A and B Tests for Rapid Detection of Influenza Virus Infection. J Clin Microbiol. 2016;54(11):2763–2766. doi:10.1128/JCM.01586-16.

